# Systems genetic analysis of binge-like eating in a C57BL/6J x DBA/2J-F2 cross

**DOI:** 10.1101/2020.06.24.168930

**Authors:** Emily J. Yao, Richard K. Babbs, Julia C. Kelliher, Kimberly P. Luttik, M. Imad Damaj, Megan K. Mulligan, Camron D. Bryant

## Abstract

**Objective:** Binge eating is a heritable quantitative trait associated with eating disorders (**ED**) and refers to the rapid consumption of a large quantity of energy-dense food that is associated with loss of control, anxiety, and depression. Binge Eating Disorder is the most common ED in adults in the US; however, the genetic basis is unknown. We previously identified robust mouse inbred strain differences between C57BL/6J and DBA/2J in binge-like eating (**BLE**) of sweetened palatable food **(PF**) in an intermittent access, conditioned place preference paradigm.

**Methods:** To map the genetic basis of BLE, we phenotyped and genotyped 128 C57BL/6J x DBA/2J-F2 mice.

**Results:** We identified a quantitative trait locus (**QTL**) on chromosome 13 influencing progressive changes in body weight across training days (LOD = 5.5; 26-39 cM). We also identified two sex-combined QTLs influencing PF intake on chromosome 5 (LOD = 5.6; 1.5-LOD interval = 21-28 cM) and 6 (LOD = 5.3; 1.5-LOD interval = 50-59 cM). Furthermore, sex-specific analyses revealed that the chromosome 6 locus was driven by males (1.5-LOD interval: 52-59 cM) and identified a female-selective QTL for BLE on chromosome 18 (LOD = 4.1; 1.5-LOD interval: 23-35 cM). Systems genetic analysis of the chromosome 6 locus for BLE using GeneNetwork legacy trait datasets from BXD recombinant inbred strains identified *Adipor2* and *Plxnd1* as two positional, functional, biological candidate genes.

**Discussion:** We identified genetic loci influencing BLE. Future studies will phenotype BXD recombinant inbred strains to fine map loci and support candidate gene nomination and validation.

## INTRODUCTION

Binge eating (**BE**) is a heritable complex trait present within the spectrum of eating disorders (**ED**), including Binge Eating Disorder (**BED**), Bulimia Nervosa (**BN**), and Anorexia Nervosa (**AN**). BE is defined by repeated bouts of ingesting large quantities of food intake over a short time period (typically less than two hours) that is associated with a loss of control, anxiety, guilt, remorse, and depression (Wolfe et al. 2009). While BE quantity, duration, and frequency are deemed important characteristics of BE (Johnson et al. 2000), the severity of loss of control [inability to eat the amount of food that was intended or planning a binge in violation of normal dietary standards (Johnson et al. 2000)] can best predict clinical impairment and psychiatric dysfunction (Vannucci et al. 2013).

BE is associated with behavioral, malnutritional, metabolic and psychiatric dysfunction, including aberrant and compensatory restrictive eating, obesity and associated health risks (da Luz et al. 2018), negative valence (Vannucci et al. 2015), mood disorders (Guerdjikova et al. 2019), and substance use disorders (Munn-Chernoff and Baker 2016). Genetic and environmental factors contribute to susceptibility to BE (Bulik, Sullivan, and Kendler 2003). Genome-wide association studies (**GWAS**) of ED have identified significant risk loci for AN, including loci near genes related to metabolic and psychiatric dysfunction (Watson et al. 2019). However, GWAS of BE, BED, and BN are lacking (Hübel et al. 2018).

We developed a binge-like eating (**BLE**) procedure in mice to measure escalation in the consumption of sweetened palatable food (**PF**) over time in an intermittent, limited access conditioned place preference (**CPP**) paradigm and subsequent compulsive-like intake in a light/dark conflict procedure (Kirkpatrick et al. 2017). We identified multiple novel genetic factors contributing to BLE. First, using a Reduced Complexity Cross (**RCC**) between closely related substrains of C57BL/6 mice (Bryant et al. 2018, 2020), we mapped a major-effect quantitative trait locus (**QTL**) near a proposed gain-of-function missense mutation in *Cyfip2* (Kumar et al. 2013) that influenced BLE (Kirkpatrick et al. 2017). *Cyfip2* +/-mice showed reduced BLE on the BLE-prone C57BL/6NJ (**B6NJ**) background (Kirkpatrick et al. 2017). As subsequent study showed that haploinsufficiency of the closely related gene *Cyfip1* also modulated BLE but in a complex manner that depended on C57BL/6 genetic background, sex, and parent-of-origin (Babbs et al. 2019). Finally, knockout mice for casein kinase 1-epsilon (*Csnk1e*), a gene whose deletion enhances behavioral sensitivity to the stimulant, rewarding, and reinforcing responses to opioids and psychostimulants (Bryant et al. 2012; Wager et al. 2014) and is associated with opioid dependence in humans (Levran et al. 2008, 2015), showed a robust, female-specific induction of BLE on the BLE-resistant C57BL/6J (**B6J**) background (Goldberg et al. 2017). These studies illustrate the utility of our BLE paradigm in identifying genetic factors exerting pleiotropic influence on addiction traits and BLE that could have clinical relevance in humans.

To expand our efforts in gene discovery of BLE, we identified a robust genetic difference in BLE between the BLE-resistant B6J inbred strain which showed very little BLE of sweetened palatable food in our intermittent, limited access CPP (Goldberg et al. 2017; Kirkpatrick et al. 2017) versus the DBA/2J (**D2J**) inbred strain which showed robust BLE (Babbs et al. 2018). In that study, we generated a small cohort of B6J x D2J-F2 mice and tested candidate loci based on the prior QTL literature regarding sweet taste (chromosome 4) and bitter taste (chromosome 6) between B6J and D2J strains (Blizard, Kotlus, and Frank 1999). We found a significant association between BLE in males and a polymorphic marker within the *Tas2r* locus on chromosome 6 (133 Mb) containing bitter taste receptors (Babbs et al. 2018) that was previously associated with variation in quinine (bitter) taste between B6J and D2J (Blizard et al. 1999; Nelson, Munger, and Boughter 2005). However, there are several remaining questions from these findings. First, is the association between *Tas2r* and BLE significant at the genome-wide level? Second, does linkage of BLE with chromosome 6 peak near*Tas2r* (∼132.5-133.5 Mb, mm10) or can it be more precisely localized? Third, is there functional evidence for candidate genes underneath this QTL that could modulate BLE? And fourth, can we identify additional genome-wide significant loci linked to BLE?

To answer these questions, we genotyped the same cohort of B6J x D2J-F2 mice (Babbs et al. 2018) genome-wide to conduct QTL analysis of BLE and determine if we could replicate candidate loci at the genome-wide level and identify new QTLs underlying BLE. Our results confirmed a major genome-wide significant, male-sensitive QTL on chromosome 6. Furthermore, we identified an additional QTL on chromosome 5 affecting BLE in both sexes and a nearly significant female-sensitive QTL for BLE on chromosome 18. We employed GeneNetwork (Chesler et al. 2004; Mulligan et al. 2017) to identify candidate genes exhibiting functional evidence for B6J/D2J polymorphisms influencing their expression and to identify the correlation of these candidate genes with other relevant behavioral and physiological traits. GeneNetwork (www.genenetwork.org/) is an online data repository and tool for analyzing thousands of historical gene expression, physiological, and behavioral traits among mouse crosses and genetic reference panels, especially in crosses and panels segregating B6J and D2J alleles (Chesler et al. 2004; Mulligan et al. 2017).

## METHODS

### Mice

All experiments were conducted in accordance with the NIH Guidelines for the Use of Laboratory Animals and were approved by the Institutional Animal Care and Use Committee at Boston University (AN-15403). Seven-week old, B6J x D2J-F_1_ mice (15 breeder pairs) were purchased from Jackson Laboratory (JAX; Bar Harbor, ME) and were habituated in the colony for one week prior to breeding in-house to generate 128 B6J x D2J-F_2_ mice for experimental testing.

### Home cage diet and experimental palatable food (PF) pellets

Chow (Teklad 18% Protein Diet, Envigo, Indianapolis, IN, USA) and tap water were provided in the home cage *ad libitum* throughout the entire study. For BLE training, sweetened PF pellets (TestDiet; 20 mg each; 5TUL diet; MO, USA) contained a metabolizable energy density of 3.4 kcal/g (21% from protein, 13% from fat, 67% from carbohydrates) were provided in an intermittent, limited access model of BLE as described below.

### Binge-like eating (BLE) and compulsive-like eating (CLE)

F2 mice were previously trained in an intermittent, limited access BLE procedure in a PF conditioned place preference (**CPP**) paradigm over 22 days (Babbs et al. 2018) as described in the **Supplementary Information** and multiple publications (Babbs et al. 2019, 2020; Kirkpatrick et al. 2017).

### Genotyping in B6J × D2J-F_2_ mice

DNA was extracted from tail snips using a salting out procedure. DNA was shipped for genome-wide genotyping on the MiniMUGA array (Neogen GeneSeek Operations, Lincoln, NE, USA). There are 3314 polymorphic SNP markers between B6J and D2J on this array that can identify parental inheritance of recombinant, chromosomal regions. Marker positions were converted from bp to sex-averaged cM prior to mapping, using the JAX Mouse Map Converter (http://cgd.jax.org/mousemapconverter).

### Data analysis

Prior to QTL analysis, F2 mice were analyzed irrespective of genotype in R using mixed model ANOVAs with Sex as a factor and Day as a repeated measure followed by unpaired t-tests (Sex comparisons) or paired t-tests (Day comparisons). Quality checking and QTL analysis were performed in R (https://www.r-project.org/) using R/bestNormalize (https://github.com/petersonR/bestNormalize) and R/qtl (Broman et al. 2003). Phenotypes were assessed for normality using the Shapiro-Wilk Test. Because the data residuals sometimes deviated significantly from normality, we used the orderNorm function to perform Ordered Quantile (**ORQ**) normalization (Peterson and Cavanaugh 2019) on all phenotypes. For sex-specific analyses, datasets were quantile-normalized separately for females and males. Genotype calls were quality checked to identify possible errors and to ensure reliable markers. We dropped any markers with a call rate of less than 95%. We used the countXO function, to examine the number of crossovers per individual and removed three outlier subjects showing more than 1000 crossovers each. Finally, we identified double-crossover genotyping errors by running the calc.errorlod function in R/qtl with an assumed genotyping error rate of 0.05. Markers with log of the odds (**LOD**) scores greater than 5 were removed. After QC, 2994 markers and phenotypes from 128 F2 samples were used in QTL analysis.

QTL analysis was performed using the “scanone” function and Haley-Knott (HK) regression. “Cohort” was included as an additive covariate and “Sex” was included as an interactive covariate in the QTL model. For separate female and male analyses, “Cohort” was included as an additive covariate. Permutation analysis (perm = 1000) was used to compute genome-wide suggestive (p < 0.63) and significance (p < 0.05) thresholds. For each significant QTL, we calculated both the Bayes credible interval and 1.5 LOD drops from the peak-associated marker. Percent phenotypic variance explained by each QTL was calculated using the “fitqtl” function.

### Identifying candidate genes and variants within the QTL intervals

The Sanger Institute Mouse Genomes Project (https://www.sanger.ac.uk/science/data/mouse-genomes-project) contains gene annotations of inbred mouse strains. We used this tool to identify polymorphic genes between C57BL/6J and DBA/2J within the male-selective chromosome 6 QTL interval (111-125 Mb) were filtered to include SNPs and insertions/deletions (indels) that were Ensembl-annotated as “high impact” (see Supplementary Information).

### Systems genetic analysis of male-selective chromosome 6 QTL for BLE in GeneNetwork

*Cis*-eQTL and PheQTL-eQTL network analysis were performed using GeneNetwork, an online data repository containing legacy SNP and transcriptome datasets and an analysis tool, to explore gene regulatory networks (Mulligan et al. 2017). Candidate genes for each QTL were identified using BXD RI data sets (**Supplementary Information**).

Sixty-seven pheQTLs with peak Likelihood Ratio Statistic (**LRS**) scores within each interval of interest were identified through the BXD Published Phenotypes dataset [BXDPublish; GN602]. PheQTL-eQTL network graphs were generated to visualize Pearson correlation coefficients greater than 0.5 or less than −0.5.

### Power analysis

Knowing that we had limited power to detect QTLs with a sample size of 128 F2 mice unless a locus was of large magnitude, we used the R package QTLdesign with the “detectable” function on D23 intake data to generate a plot representing power versus variance explained for an additive QTL (p<0.05) (Sen et al. 2007).

## RESULTS

### Changes in BW, PF intake, PF-CPP, and power analysis in B6J x D2J-F2 mice

In examining BW across BLE training (D2-D18) and CLE assessment (D23), males showed a significantly greater percent increase in BW and a greater slope of increase in BW gain compared to females (**Figure 1A,B**). Despite showing less % increase in BW, females consumed significantly *more* PF than the males on D4, D16, and D23 (**Figure 1C**); however, the slope of intake did not differ significantly (**Figure 1D:** p = 0.1). Interestingly, there was a significant negative correlation between the slope of % BW gain (D2-D18) and the slope of % BW of PF consumed (D2-18) (**Figure 1E**), indicating that greater escalation of PF intake predicted less rapid BW gain. In an intermittent access procedure, rodents learn to reduce home cage chow intake in anticipation of the more reinforcing PF (Cottone et al. 2008) which could explain this negative relationship.

**Figure 1.**
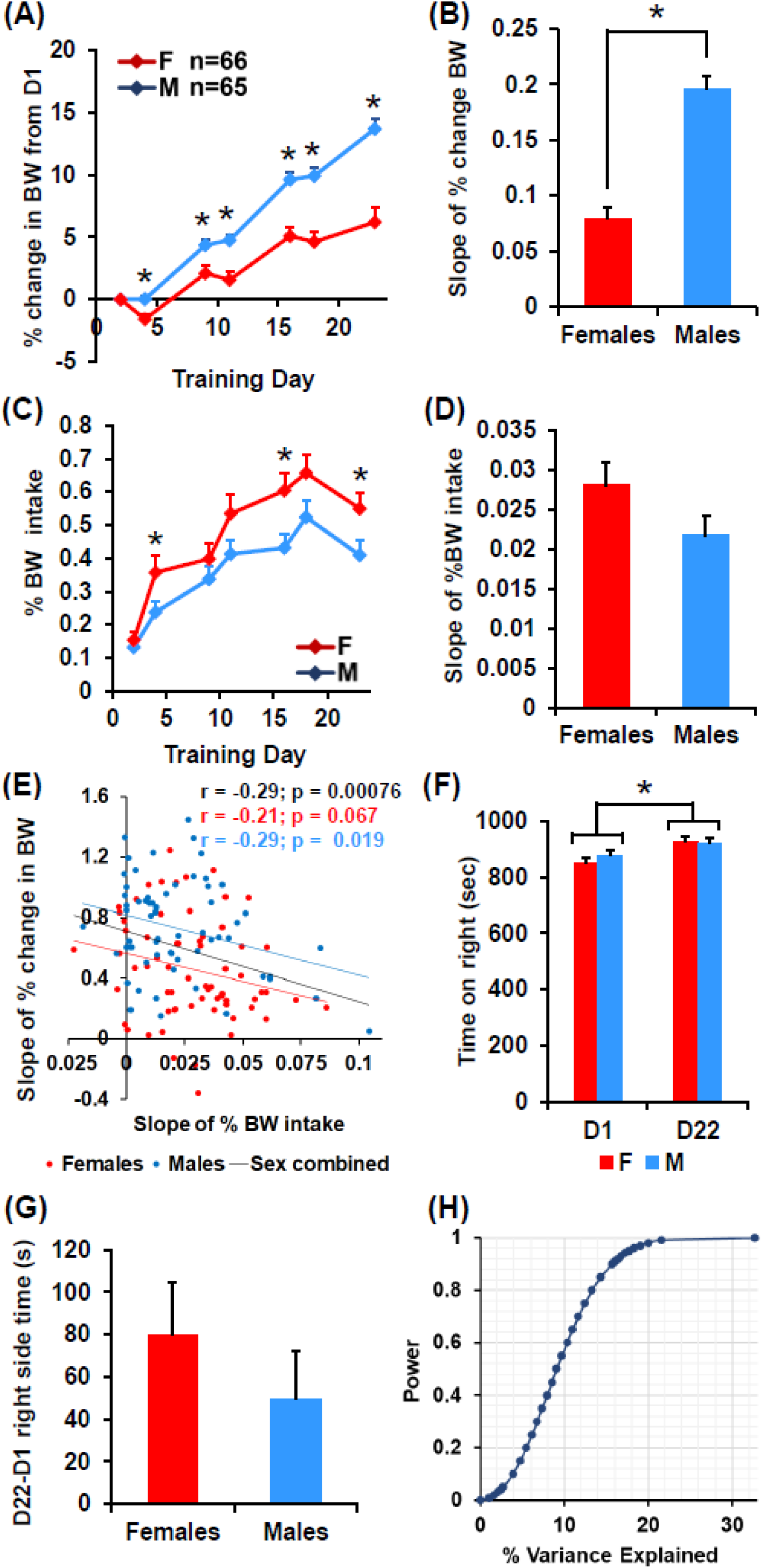
Percent change in BW, PF intake, PF-CPP, and power analysis in B6J x D2J-F2 mice. **(A):** Change in BW (% of D2) across BLE training days through D22; just prior to assessment of compulsive-like eating (**CLE**) on D23. There was a main effect of Day (F_5,645_ = 169.26, p < 2 × 10^−16^), Sex (F_1,129_ = 27.8, p = 5.5 × 10^−7^), and an interaction (F_5,645_ = 13.29, p = 2.4 × 10^−12^). On all PF days subsequent to the first day (D2) of PF exposure (i.e., D4, D9, D11, D16, D18, and D23), males showed a significantly greater % increase in BW compared to the females (**\***all p’s ≥ 0.0003). **(B):** Slope of % increase in BW from D2 through D22 in F2 females (n = 66) and in F2 males (n = 65). Unpaired Student’s t-test revealed a significantly steeper rise in % increase in BW over BLE training days in males compared to females (t_129_ = −7.69, **\***p = 3.35 × 10^−12^). **(C):** PF intake (%BW) over BLE training days in F2 females and males. There was a main effect of Day (F_6,774_ = 59.06, p < 2 × 10^−16^) and Sex (F_1,129_ = 4.64, p = 0.03), but no interaction (p = 0.12). Females showed greater intake on D4, D16, and D23 (t_129_ = 2.0, 2.54, 2.18; *p’s = 0.048, 0.012, 0.031). **(D):** Analysis of slope of PF intake (%BW) in F2 females and F2 males indicated no significant difference (p = 0.10). **(E):** Negative correlation between the slope of PF intake (%BW) and the slope of weight gain (% of D2) for the sex-combined (black: r = −0.29, t_129_ = 3.45; p = 00076), female (red: r = −0.23; t_129_ = 1.87; p = 0.067), and male (blue: r = −0.29; t_129_ = 2.41; p = 0.019) datasets. **(F):** Time spent on the PF-paired side on D1 prior to BLE training and on D22 post-BLE training. There was a main effect of Day (F_1,129_ = 14.66, *p = 2 × 10^−4^) which indicated significant PF-CPP, but no effect of Sex (p = 0.619), and no interaction (p = 0.37). **(G):** Analysis of the difference in time spent on the PF-paired side between D1 and D22 (D22-D1, s) in females and males confirmed no significant difference (t_129_ < 1). **(H):** Power versus effect size (% variance explained) for an additive QTL model and a sample size of 128 F2 mice (p < 0.05). 0.2, 0.4, 0.6, and 0.8 power is achieved with an observed effect size of 5.45%, 7.93%, 10.31%, and 13.32% variance explained, respectively.

In examining PF-CPP, there was a main effect of Day (**Figure 1F: \***p = 2.0 × 10^−4^) but no interaction with Sex, indicating significant PF-CPP, regardless of Sex. Analysis of the change in preference (D22-D1) confirmed no significant sex difference (**Figure 1G**).

**Figure 1H** shows power versus effect size (% variance explained) for an additive QTL with our sample size (128 F2 mice; p < 0.05). Twenty, 40, 60, and 80% power can be achieved with an effect size of 5.45%, 7.93%, 10.31%, and 13.32% phenotypic variance explained, respectively.

### Identification of QTLs underlying differences in BW and PF intake but not PF-CPP

**Table 1** lists the details of the QTLs discussed below. **Supplementary Figure 1** provides a visual heat map of the QTLs via R/qtlcharts (Broman 2015). We identified a genome-wide significant QTL on chromosome 13 for D23 BW that explained 64% of the variance (**Figure 2A**). There was a day-dependent increase in linkage that was significant by D23 (**Figure 2B**). Sex-specific analyses revealed that only females showed a significant peak that was more distally located than the nonsignificant males-only peak (**Figure 2C**). The effect plots of the peak loci show an increase in BW associated with the D2J allele (**Figure 2D-F**).

**Table 1.**

| QTLs for body weight (BW) and PF intake |  |  |  |  |  |  |
| --- | --- | --- | --- | --- | --- | --- |
| Trait | Chr. | Peak, cM (Mb) | LOD | 1.5 LOD interval, cM (Mb) | Bayes interval, cM (Mb) | % variance explained |
| D23 BW | 13 | 31 cM (58 Mb) | 5.5 | 26-39 cM (46-73 Mb) | 27-38 cM (53-72 Mb) | 64% |
| D2 intake | 5 | 26 cM (46 Mb) | 5.8 | 21-28 cM (40-53 Mb) | 13-61 cM (28-121 Mb) | 19% |
| D23 intake | 6 | 53 cM (114 Mb) | 5.4 | 50-59 cM (110-125 Mb) | 42-72 cM (93-141 Mb) | 21% |
| D2 intake (M) | 5 | 25 cM (45 Mb) | 4.4 | 12-66 cM (25-128 Mb) | 8-66 cM (17-128 Mb) | 16% |
| D18 intake (M) | 5 | 11 cM (24 Mb) | 4.2 | 7-25 cM (16-46 Mb) | 6-25 cM (15-45 Mb) | 24% |
| D16 intake (M) | 6 | 53 cM (114 Mb) | 4.3 | 43-60 cM (94-126 Mb) | 30-67 cM (65-136 Mb) | 23% |
| D23 intake (M) | 6 | 53 cM (114 Mb) | 5.9 | 52-59 cM (111-125 Mb) | 47-72 cM (100-141 Mb) | 32% |
| Slope (F) | 18 | 24 cM (45 Mb) | 4.1 | 23-35 cM (43-62 Mb) | 23-33 cM (43-58 Mb) | 23% |

D = Day of protocol; F = females only analysis; M = males-only analysis; cM = centimorgans; Mb = megabases

QTL analysis of sex-combined PF intake revealed genome-wide significant QTLs on chromosomes 5 and 6 for D2 PF intake and D23 PF intake, respectively (**Figure 3A**). For chromosome 5, the effect plot of the peak locus showed an increase in D2 PF intake with increasing D2J alleles (**Figure 3B,C**). For chromosome 6, the D2J allele was also associated with increased intake (**Figure 3D,E**). The chromosome 6 QTL clearly showed a progressive increase in linkage across PF intake assessment days (**Figure 3D**), indicating that the strength of genetic linkage reflects the strength of increased PF intake.

**Figure 2.**
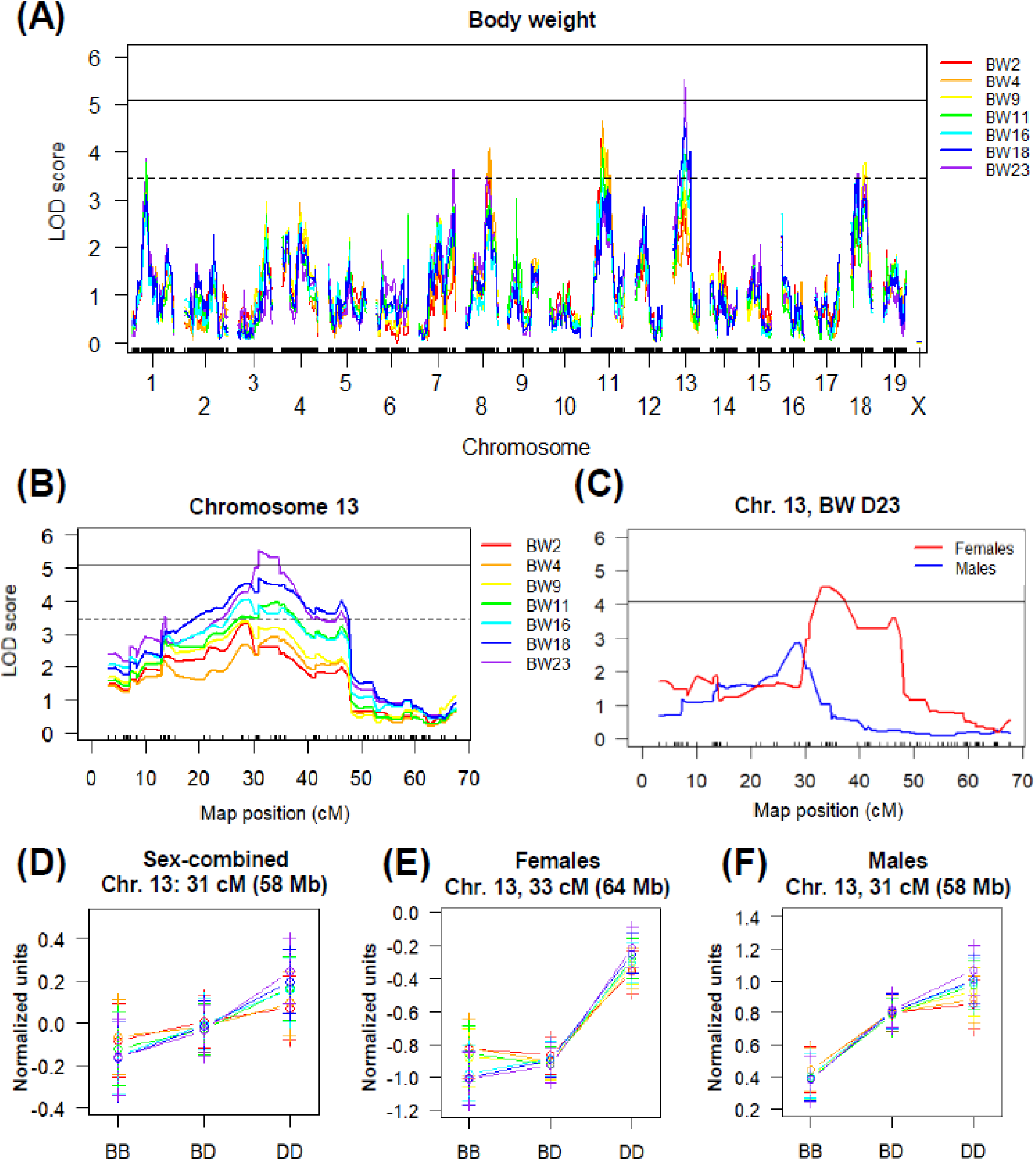
Emergence of a genome-wide significant QTL on medial chromosome 13 influencing BW during BLE and CLE. **(A):** Genome-wide QTL plot for BW across assessment days for PF intake revealed a significant QTL on chromosome 13 for Day 23 BW [LOD = 5.54, peak = 30.79 cM (57.85 Mb), Bayes C.I.: 52.51-71.71 Mb, 1.5 LOD: 51.46-72.52 Mb]. The numbers indicate the chromosome-#. The solid horizontal line for panels A-C indicates significance threshold (p < 0.05) and the dotted horizontal line indicates the suggestive threshold (p < 0.63). 64% of the phenotypic variance was explained by the chromosome 13 QTL. **(B):** Chromosome 13 QTL plot of BW for each day of assessment of PF intake. The rainbow color scheme illustrates a clear day-dependent increase in linkage across days/spectrum (red -> purple) that is significant by the final day of PF intake assessment on D23 (purple trace) prior to CLE assessment in the light/dark box. **(C):** Female and male chromosome 13 QTL plots for BW on D23. Females showed a more distal peak on chromosome 13 (red trace; peak = 32.98 cM, 63.76 Mb) compared to the males (blue trace; peak = 30.79 cM, 57.85 Mb) that drove the overall sex-combined QTL signal. **(D):** Sex-combined effect plot at the peak locus for D23 BW (chromosome 13: 30.79 cM, 57.85 Mb). There is an increase in quantile-normalized BW with increasing D2J (D) alleles. **(E):** Female effect plot at the peak chromosome 13 locus for D23 BW (32.98 cM, 63.76 Mb) shows an increase in quantile-normalized BW with increasing D alleles. **(F):** Male effect plot of the peak locus on chromosome 13 for D23 BW (30.79 cM, 57.85 Mb) shows an increase in quantile-normalized BW with each D allele.

**Figure 3.**
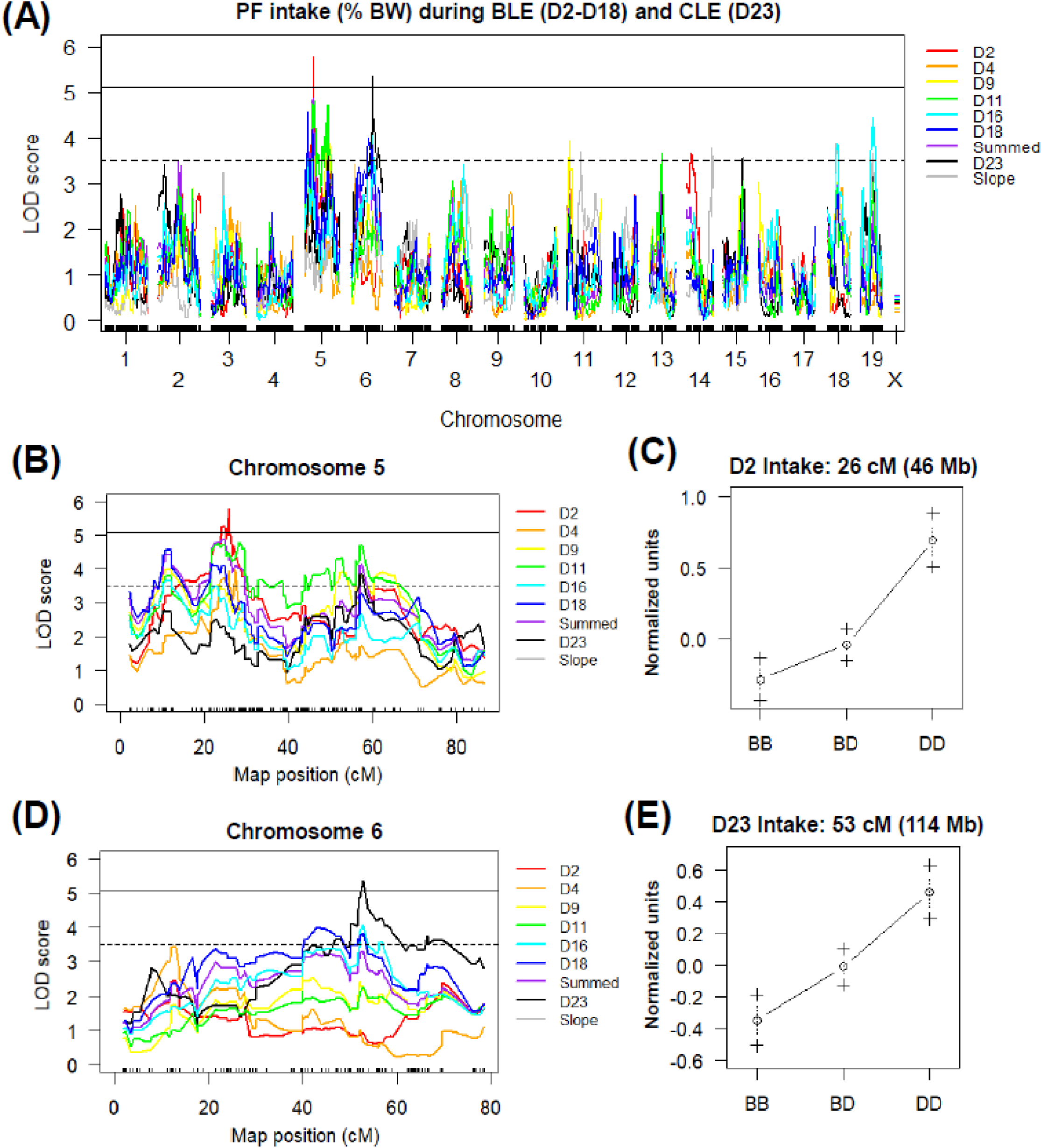
Sex-combined QTLs influencing PF intake on chromosomes 5 and 6. **(A):** Genome-wide QTL plot revealed significant QTLs for PF intake (% BW consumed) on chromosomes 5 and 6. The chromosome 5 QTL was significant for D2 intake [LOD = 5.77, peak = 25.82 cM (46.20 Mb), Bayes C.I.: 27.67-121.03 Mb, 1.5 LOD: 40.22-52.51 Mb] and explained 19% of the phenotypic variance. The chromosome 6 QTL was significant for D23 intake [LOD = 5.36, peak = 52.86 cM (113.80 Mb), Bayes C.I.: 92.99-141.19 Mb, 1.5 LOD: 109.61-124.76 Mb] and explained 21% of the phenotypic variance. The solid horizontal line for panels A, B, and D indicates significance threshold (p < 0.05) and the dotted horizontal line indicates the suggestive threshold (p < 0.63). **(B):** The chromosome 5 QTL plot shows a significant QTL for PF intake on D2 (red trace). **(C):** The effect plot of the peak chromosome 5 locus (25.82 cM, 46.20 Mb) shows an increase in quantile-normalized D2 intake with each copy of the D2J (D) allele. BB = homozygous for B6J allele; BD = heterozygous; DD = homozygous for D2J allele. **(D):** The chromosome 6 QTL plot shows a significant QTL for D23 PF intake in the light/dark box. **(E):** The effect plot of the peak locus on chromosome 6 (52.86 cM, 113.80 Mb) shows an increase in quantile-normalized D23 intake with each copy of the D allele.

Because we identified sex-dependent candidate loci for BLE (Babbs et al. 2018), we conducted separate QTL analyses for females and males. For males, we identified QTLs on chromosomes 5 and 6 that mirrored the sex-collapsed results (**Figure 4A**). For chromosome 5, a significant QTL was again identified for D2 intake (**Figure 4B**) and a second, more proximal chromosome 5 QTL for D18 intake (**Figure 4B**). The D2J allele was associated with increasd PF intake (**Figure 4C**).

**Figure 4.**
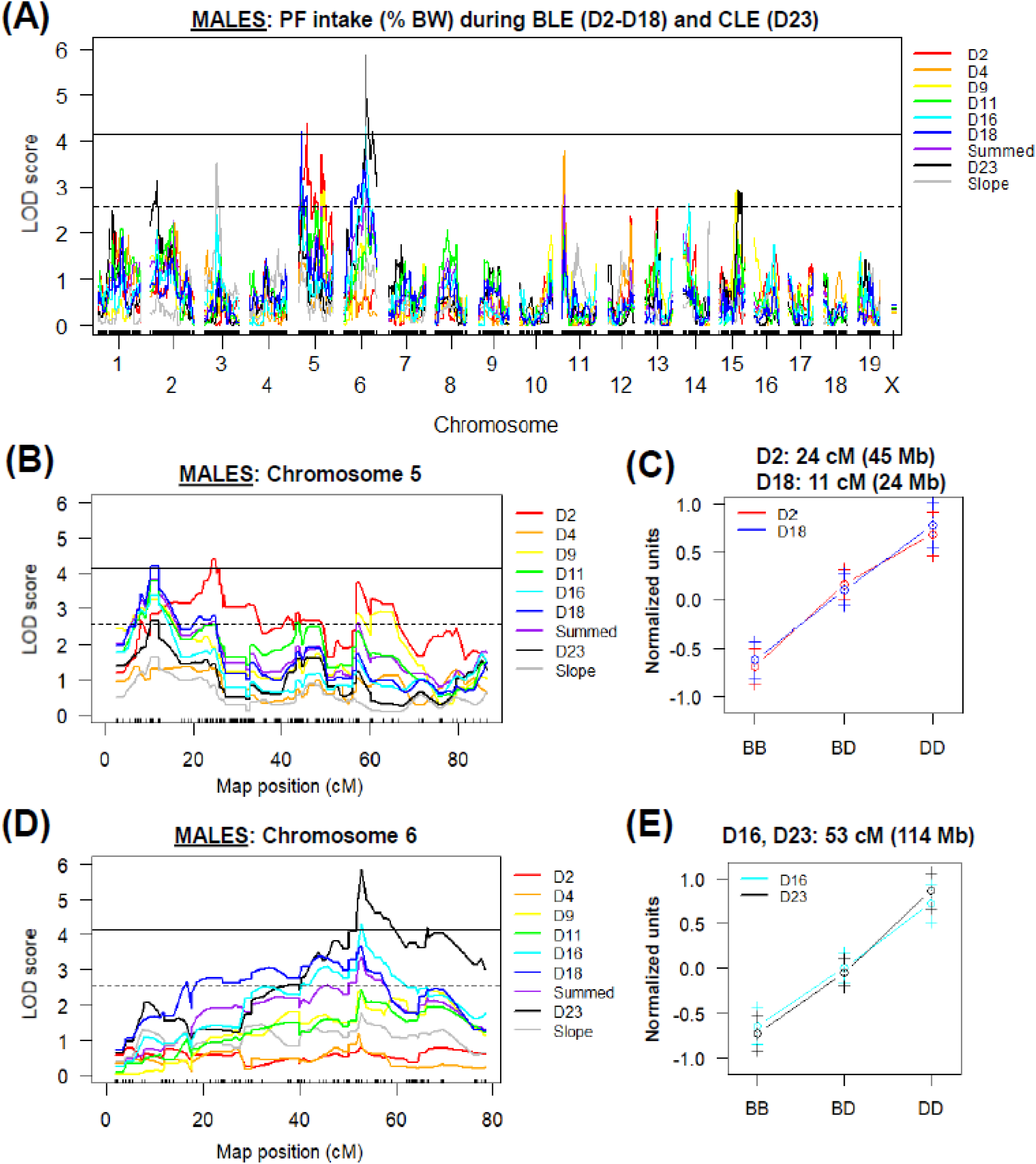
Genome-wide QTL analysis of PF intake in F2 males. **(A):** Genome-wide significant QTLs were identified for males on chromosomes 5 and 6. Chromosome 5 contained QTLs for D2 intake [LOD = 4.22, peak = 24.45 cM (44.78 Mb), Bayes C.I.: 17.13-141.23 Mb, 1.5 LOD: 24.80-127.95 Mb, 16% variance explained] and for D18 intake [LOD = 4.26, peak = 10.66 cM (24.07 Mb), Bayes C.I.: 13.73-45.06 Mb, 1.5 LOD: 15.88-45.58 Mb, 24% variance explained]. Chromosome 6 contained QTLs for D16 intake [LOD = 4.29, peak = 52.86 cM (113.80 Mb), Bayes C.I.: 65.01-138.58 Mb, 1.5 LOD: 94.06-125.62 Mb, 23% variance explained] and for D23 intake [LOD = 5.74, peak = 52.86 cM (113.80 Mb), Bayes C.I.: 100.46-141.19 Mb, 1.5 LOD: 111.06-123.63 Mb, 32% variance explained]. The solid horizontal line for panels A, B, and D indicates the significance threshold (p < 0.05) and the dotted horizontal line indicates the suggestive threshold (p < 0.63). **(B):** The QTL plot for chromosome 5 shows significant QTLs for D2 intake (red trace) and D18 intake (blue trace). **(C):** The effect plots for chromosome 5 at the peak loci for D2 intake (24.45 cM, 44.78 Mb) and D18 intake (10.66 cM, 24.07 Mb) show increased normalized PF intake with increasing D alleles. **(D):** The chromosome 6 QTL plot shows a significant QTL for PF intake on D16 (light blue trace) and D23 (black trace). **(E):** Effect plots for the chromosome 6 QTLs at the peak locus (52.86 cM, 113.80 Mb) shows an increasing effect of the D allele on PF intake for D16 and D23.

A significant chromosome 6 QTL was also identified for D16 intake in males (**Figure 4D**) and for D23 intake during CLE (**Figure 4D**). The D2J allele was associated with increased PF intake (**Figure 4E**).

For females, there was a nearly significant QTL on chr. 18 (p = 0.052) underlying the slope of escalation in PF intake (**Figure 5A,B**). Interestingly, the effect plot of the peak locus suggested an overdominance effect, with heterozygotes showing the greatest normalized escalation compared to homozygous genotypes (**Figure 5C**).

**Figure 5.**
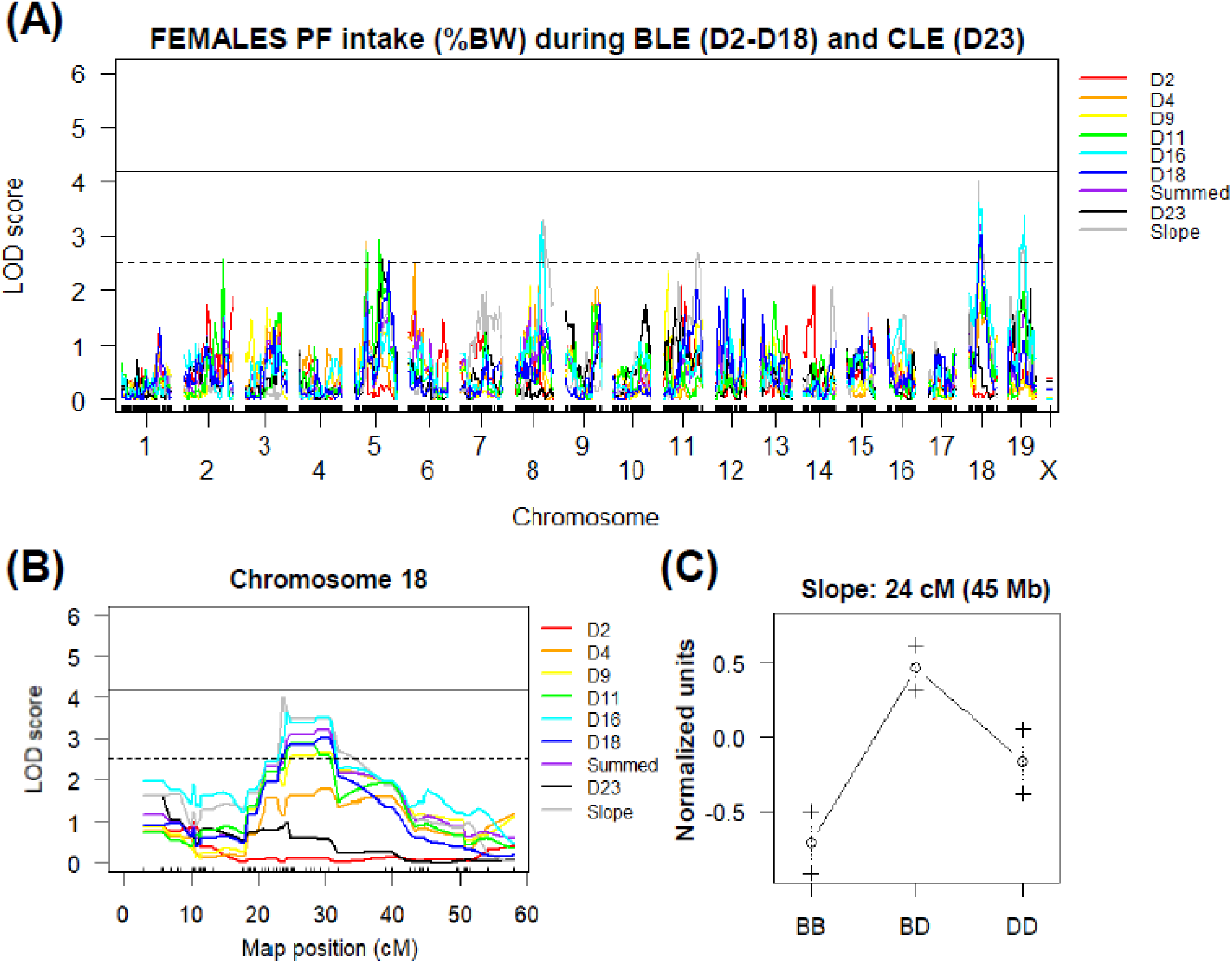
Genome-wide QTL analysis of PF intake in F2 females. **(A):** There was one, nearly significant QTL on chromosome 18 in females (p = 0.052) for the slope of intake across PF training days [LOD = 4.11, peak = 23.81 cM (44.88 Mb), Bayes C.I.: 43.16-58.40, 1.5 LOD: 42.74-61.80 Mb]. The chromosome 18 QTL explained 23% of the variance in the slope of PF intake across days. Solid horizontal line for panels A and B indicates significance threshold (p < 0.05), dotted horizontal line indicates suggestive threshold (p < 0.63). **(B):** Chromosome 18 QTL plot shows a peak on the medial portion. **(C):** Chromosome 18 effect plot at the peak locus (23.81 cM, 44.88 Mb) shows evidence for an overdominance effect whereby the heterozygous BD genotype displays the greatest normalized slope value.

For QTL analysis of PF-CPP, there were no genome-wide significant QTLs for the D22-D1 right side time measure (**Supplementary Figure 2**).

To summarize, the results indicate that the significant QTLs detected with our sex-collapsed data, in particular the QTL on chr. 6, were driven primarily by males, while the suggestive peaks on chromosomes 18 and 19 were driven by females.

### eQTL and PheQTL-eQTL networks identify candidate genes for the chromosome 6 QTL for D23 PF intake in males (111-125 Mb)

To identify positional candidate genes for the male-selective chromosome 6 locus influencing PF intake on D23, we employed the bioinformatic pipeline illustrated in **Figure 6**. The chromosome 6 QTL was prioritized because it was highly significant, narrow in size, and showed a progressive, day-dependent increase in the strength of genetic linkage with PF intake. Using the Sanger Institute Mouse Genomes Project (https://www.sanger.ac.uk), we generated a gene list containing 57 polymorphic, protein-coding genes with high-impact variant annotations within the 1.5 LOD confidence interval (chromosome 6: 111 Mb-125 Mb; **Supplementary Table 1**). Next, we used GeneNetwork (Mulligan et al. 2017) to identify eQTLs within BXD RI gene expression datasets that were associated with these genes in several brain regions (**Supplementary Information**) and then filtered genes to those possessing a maximum LRS score within our QTL interval. We then examined correlations among genes containing eQTLs and published behavioral and physiological phenotypes with QTLs containing peak LRS scores within the male-selective chromosome 6 locus. Using the BXDPublish dataset on GeneNetwork, we identified 67 traits with QTL peaks located within the chromosome 6 QTL interval; 18 of these traits were included in the analysis (**Supplementary Table 2**).

**Figure 6.**
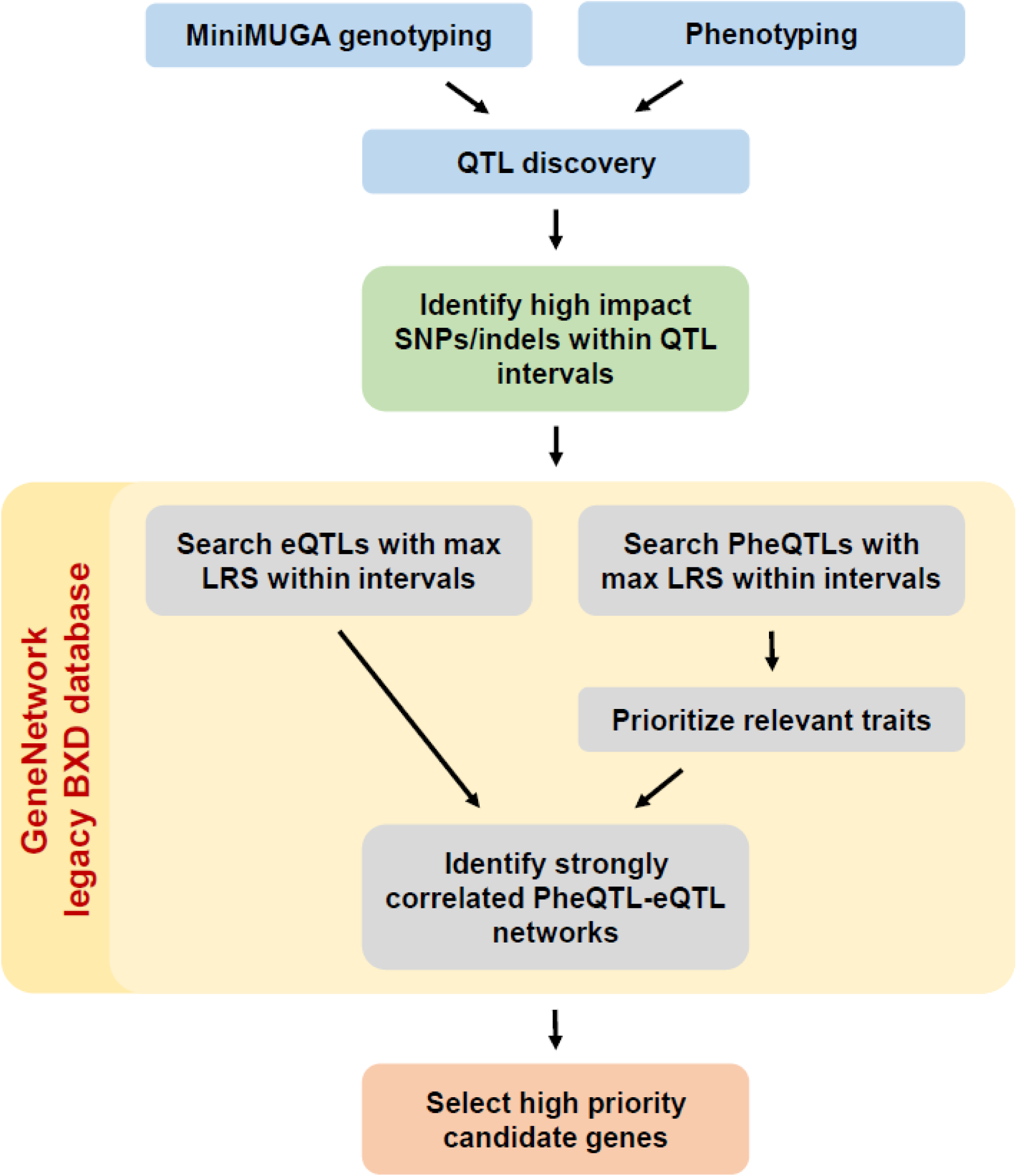
Schematic of bioinformatics pipeline for identifying positional, functional candidate genes for PF intake and associated phenotypes.

To illustrate the strongest PheQTL-eQTL correlations, we generated a graph with all genes in each dataset and all of our selected phenotype records. Three genes (*Adipor2, Plxnd1*, and *Rad18*) were strongly correlated with at least two traits. Thus, network graphs for each of these three genes were generated in GeneNetwork to further examine the correlations among phenotypes. *Cis*-eQTLs were identified for *Adipor2* in nucleus accumbens and ventral tegmental area and eight directly connected nodes were identified between gene and phenotype (**Figure 7**). *Cis*-eQTLs were identified for *Plxnd1* in hypothalamus, amygdala, and striatum and eight connected nodes (**Supplementary Figure 3**). *Cis*-eQTLs were identified for *Rad18* in prefrontal cortex, nucleus accumbens, striatum, and ventral tegmental area and 10 connected nodes (**Supplementary Figure 4**). Thus, *Adipor2, Plxnd1*, and *Rad18* are three high priority candidate genes based on the functional evidence at the level of DNA sequence, gene expression, and in turn, the connectivity of differential expression of these genes with other phenotypes relevant to PF intake.

**Figure 7.**
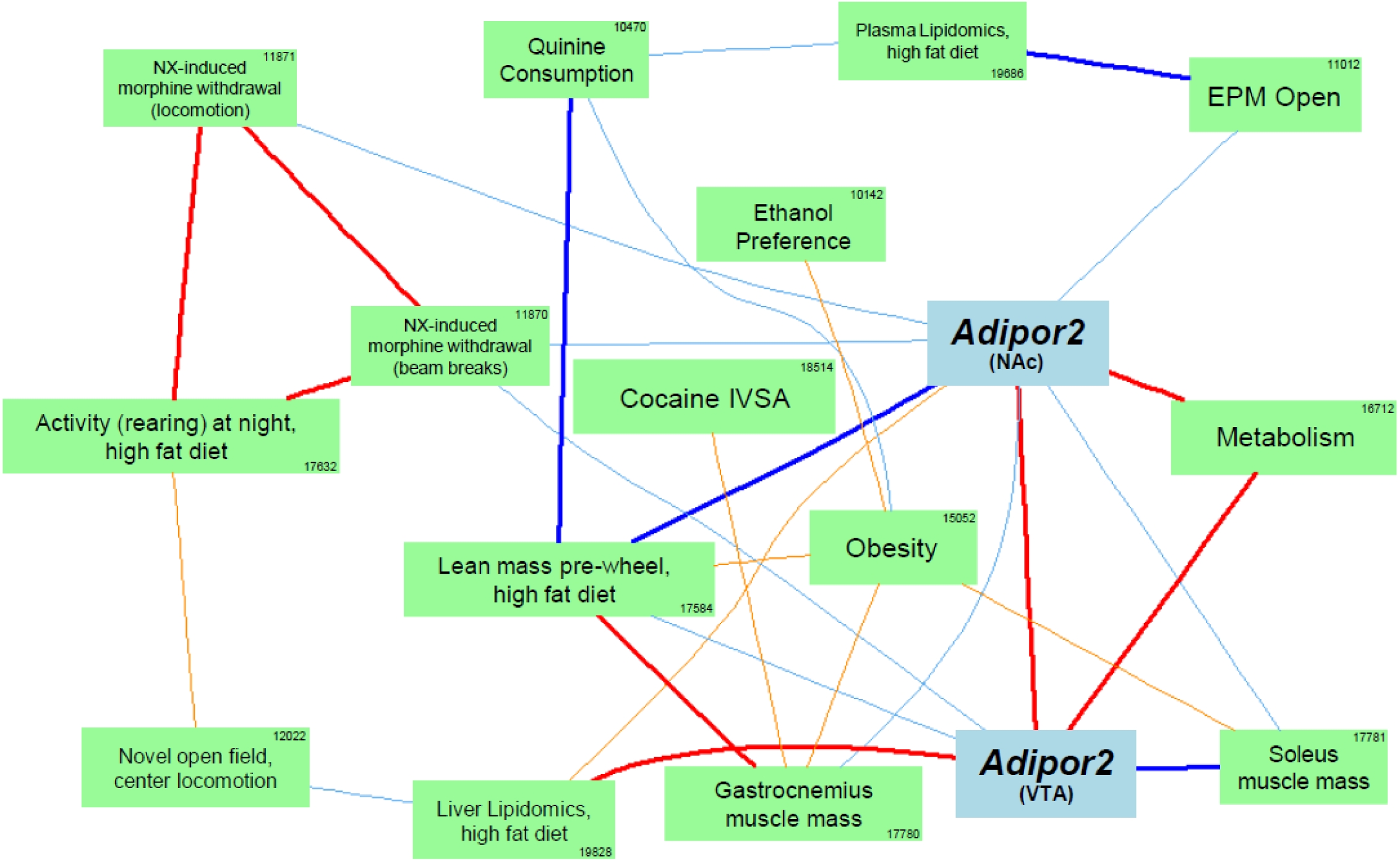
PheQTL-eQTL network graph generated via GeneNetwork for *Adipor2* within the male-specific chromosome 6 QTL interval (111-125 Mb) shows *cis-*eQTLs in two brain regions (nucleus accumbens and ventral tegmental area) with eight phenotypic nodes. Correlated PheQTLs include traits involved in metabolism, lipidomics, body mass, anxiety, and morphine withdrawal (**Supplementary Table 2**). Bold blue lines indicate Pearson correlation coefficients of −1 to −0.7, normal light blue: −0.7 to −0.5, bold red: 0.7 to 1, normal orange: 0.5 to 0.7. The number inside each green box indicates the trait’s BXDPublish Record ID number (see Supplementary Table 2). Only connected nodes are shown.

## DISCUSSION

We identified major QTLs during BLE and CLE (**Table 1**), including a chromosome 13 QTL for BW gain (**Figure 2**), a chromosome 5 locus for initial PF intake (**Figure 3B,C**), a chromosome 6 locus (**Figure 3D,E**) that was male-selective (**Figure 4D,E**) and influenced final PF intake, and a nearly significant, female-selective chromosome 18 QTL for the slope in escalation of PF intake (**Figure 5**). The BW QTL was separate from loci influencing PF intake, indicating a genetic dissociation. In fact, the slope in escalation of PF intake during BLE was somewhat negatively correlated with the slope of BW gain (**Figure 1E**), suggesting a learned, reduced intake of the less reinforcing home cage chow in BLE-prone animals in anticipation of the more reinforcing PF (Cottone et al. 2008).

We resolved the male-selective chromosome 6 locus for D23 PF intake (Babbs et al. 2018) to a region peaking at 114 Mb and spanning 111-125 Mb (**Table 1; Figure 4D**). The*Tas2r* locus containing bitter taste recpetors (132.5-133.5 Mb) lies outside of this 1.5-LOD support interval near the distal edge of the Bayes interval (100-141 Mb), 24 Mb distal from the peak. Although these observations do not rule out the *Tas2r* cluster as a source of the QTL, they do call into question whether *Tas2r* is the primary contributor to differences in PF intake. The original QTL for bitter (quinine) taste sensitivity using the same F2 cross was more distal and peaked much closer to *Tas2r* at 129 Mb (62 cM; D6Mit338) and spanned 112-139 Mb (49-67 cM: D6Mit287-D6Mit198) (Blizard et al. 1999). Other studies identified the same locus for bitter taste sensitivity in other crosses with C57BL/6J, including sucrose octaacetate with NZB/BINJ spanning 87 Mb (38 cM: D6Mit9) to 146 Mb (78 cM: D6Mit14) (Le Roy, Pager, and Roubertoux 1999). Subsequent analysis of quinine sensitivity in the BXD recombinant inbred (**RI**) strain panel (comprising fixed alleles from B6J and D2J) resolved the locus to a sharp peak squarely flanking *Tas2r* (D6Mit13; 132.6 Mb), spanning 125.4 Mb (D6Mit254) to 134.2 Mb (D6Mit374) (Nelson et al. 2005). High resolution mapping confirmed the same peak marker and interval for sucrose octaacetate taste aversion (Bachmanov et al. 2001). To summarize, we located a more proximal chromosome 6 peak and locus for BLE compared to the historical bitter taste locus, suggesting additional genetic factors besides the *Tas2r* locus contribute to variance in BLE.

What are the causal genetic factor(s) upstream of *Tas2r* on chromosome 6 that underlie BLE in males? To identify positional candidate genes based on functional evidence and correlations with historical phenotypes, systems genetic analysis using legacy BXD RI datasets from GeneNetwork (Mulligan et al. 2017) identified *Adipor2* (adiponectin receptor 2) as a top candidate gene, which codes for a seven transmembrane domain cell-surface receptor for the protein adiponectin, an adipokine secreted by white adipose tissue that regulates the metabolism of lipids and glucose and insulin sensitivity (Yamauchi et al. 2014). Adiponectin acts on *Adipor2* and *Adipor1* in the periphery and in the brain (hypothalamus, brainstem, pituitary, cortex) to regulate energy homeostasis and other processes such as synaptic plasticity and neurogenesis by signaling through AMPK, p38 MAPK, JNK, PPARα, and NF-kB (Bloemer et al. 2018; Thundyil et al. 2012; Yamauchi et al. 2007). There are 212 variants within *Adipor2* that distinguish the D2J strain from the B6J strain (https://www.sanger.ac.uk), including one 5’ UTR SNP, three, 3’ UTR SNPs, and over 60 nmd intronic variants (see “high impact variants” in **Supplementary Table 1**). Serum levels of adiponectin are inversely correlated with BMI and risk for diabetes and are decreased in patients with BED and increased in patients with AN (Khalil and El Hachem 2014). *ADIPOR2* is implicated in diabetes, obesity, high-fat feeding, and metabolism (Yamauchi et al. 2014). *Adipor2* knockout mice show lower body fat and increased resistance to high-fat diet-induced obesity, along with improved glucose tolerance, increased locomotor activity, and energy metabolism whereas *Adipor1* knockouts showed largely opposite phenotypes (Bjursell et al. 2007). In addition, high-fat feeding is associated with lower adiponectin and higher levels of both *Adipor1* and *2* (Bullen et al. 2007). Interestingly, adult males (mice and humans) show lower plasma adiponectin than females (Arita et al. 1999; Gui, Silha, and Murphy 2004), providing evidence that sex differences in the adiponectin system could underlie male-selective genetic effects of the chromosome 6 locus containing *Adipor2* on BLE (**Figures 3-4**).

Although *Adipor2* is an interesting candidate gene, our BLE regimen is relatively abbreviated and is unlikely to induce much metabolic dysfunction as evidenced by a lack of increased weight gain relative to control chow pellet training (Babbs et al. 2018), a decreased correlation between slope of PF intake and BW gain (**Figure1E**), and a genetic dissociation between QTLs for BW (**Figure 2**) and BLE (**Figures 3-5**). Therefore, could *Adipor2* dysfunction contribute to earlier physiological processes that initiate progression to BLE? *Adipor1, Adipor2*, and T-cadherin transcripts (third adiponectin receptor) can be detected in taste receptor cells (Crosson et al. 2019), suggesting that saliva-derived adiponectin could impact taste processing to influence eating. Also, adiponectin-induced activation of *Adipor1* expressed on dopamine neurons in the VTA decreased spontaneous neuronal activity and firing and reversed stress-induced increase in dopamine neuron firing and anxiety-like behavior and *Adipor1* haploinsufficiency increased dopamine neuron firing and anxiety-like behavior (Sun et al. 2019). These findings suggest that the adiponectin system, traditionally thought to regulate metabolic/homeostatic functions, could communicate taste information (e.g., hedonic versus aversive) to the mesolimbic dopaminergic reward system to influence development of BLE.

A second candidate gene based on GeneNetwork analyses was *Plxnd1* (**Supplementary Figure 3**) which codes for plexin D1, a cell surface receptor for class 3 semaphorins that regulates migration of cell types, including neuronal axon guidance and synapse formation [e.g., in striatum (Ding et al. 2011)] and vascular development (Oh and Gu 2013). There are 119 variants in *Plxnd1* distinguishing D2J from B6J (https://www.sanger.ac.uk), including three, 3’ UTR variants, two splice site variants, one 5’ UTR variant, and one missense variant (see “high impact variants” in **Supplementary Table 1**). Semaphorin 3E/plexin D1 also mediates macrophage recruitment to visceral adipose tissue during obesity to promote cytokine expression, inflammation and insulin resistance (Schmidt and Moore 2013). PLXND1 has been associated with body fat distribution in humans (Justice et al. 2019) and a nominal genetic association was identified with lipolysis (Strawbridge et al. 2016) – the hydrolysis of lipids to fatty acids in adipocytes that contributes to metabolic dysfunction and obesity. *Plxnd1* function is required for normal adipocyte morphology and number, body fat distribution, and insulin sensitivity (Minchin et al. 2015). We observed some interesting correlations of *Plxdn1* expression with addiction traits such as cocaine reinforcement, ethanol preference, and morphine withdrawal as well as behavioral models for psychiatric traits such as anxiety-like behavior and novelty seeking (**Supplementary Figure 3**). Self-administration of the mu opioid receptor agonist oxycodone was associated with an upregulation of *Plxnd1* in the nucleus accumbens (Yuferov et al. 2018). Thus, *Plxnd1* is a reasonably strong, second candidate gene within the chromosome 6 locus that could underlie male-selective differences in PF intake during BLE.

A third candidate gene based on GeneNetwork analyses was *Rad18* (**Supplementary Figure 4**) which is located at 112.62 Mb. *Rad18* codes for RAD18 E3 ubiquitin protein ligase which is a protein that is part of the DNA repair pathway and associates with other Rad proteins and ubiquitinating proteins following DNA damage and is involved in recombination (Ting, Jun, and Junjie 2010). There are 95 *Rad18* variants distinguishing B6J from D2J https://www.sanger.ac.uk), including five, 3’ UTR variants, 25 intronic nmd variants, and one 7 Kb structural variant (deletion) (see **Supplementary Table 1** for high impact variants). *Rad18* dysfunction can lead to mutagenesis, carcinogenesis, and tumorigenesis (Yang et al. 2018). Expression QTLs from several brain tissues were linked to *Rad18* expression which, in turn, were correlated with several traits related to BW, metabolism, emotional, and substance use disorder traits (**Supplementary Figure 4**). However, to our knowledge, there is no known function of *RAD18* in eating behavior, eating disorders, obesity, metabolic function, or psychiatric disorders. Thus, although there is positional and functional evidence to support *Rad18* as a candidate gene for BLE, there is very little, if any evidence from the literature.

There are some limitations to this study. First, our sample size was only powered to detect QTLs of large magnitude (> 13%; **Figure 1H**). A larger F2 sample size will permit detection of smaller-effect QTLs and explain additional variance. Second, QTL resolution in F2 mice is notoriously poor. To overcome this limitation, careful selection of a subset of BXD-RI strains will allow us to fine-map the QTLs reported here. For example, there are 54 BXD-RI strains containing at least one historical recombination event within the 111-125 Mb interval on chromosome 6 (https://www.genenetwork.org). Another limitation is that we limited our bioinformatics exercise to genes containing Ensembl-defined high impact variants that were associated with eQTLs in multiple brain tissues. Causal variants could lie within other genes that lack polymorphisms with “high impact” designation, within genes that change protein function without modulating transcript levels, or within intergenic regions not assigned to any nearby genes. Second, we only assessed consumption of one particular PF diet that was essentially sweetened chow. Although the current dataset cannot speak to the specificity of the observed QTLs for the sweetened component of the PF, note that our prior study found very little parental strain differences in control chow pellet intake between B6J and D2J (Babbs et al. 2018).

In summary, we demonstrated a genome-wide significant, male-sensitive QTL on chromosome 6 that influenced BLE and identified at least two positional, functional candidate genes that have supportive biological evidence from the literature, including *Adipor2* and *Plxnd1*. We also identified a novel chromosome 5 QTL influencing BLE in both sexes. Finally, we identified a nearly significant, female-sensitive locus on chromosome 18 influencing BLE. Phenotyping and fine mapping in a population with more recombination events like the BXD RI panel will reduce the number of candidate genes and variants. Gene/variant editing and validation will permit the study of gene function in the context of multiple eating disorder models, including additional diets (e.g., high fat diet), regimens (cycles of food restriction and binge eating, stress) and comorbidity with other ED models (e.g., activity-based anorexia) and other psychiatric disorders (mood, substance use).

## ACKNOWLEDGEMENTS

This work was funded by R21DA038738 (C.D.B.), R01DA039168 (C.D.B.), and R01CA221260 (M.I.D., C.D.B.)

## AUTHORSHIP STATEMENT

E.J.Y. conducted a majority of the statistical analyses and data presentation and contributed to the writing of the manuscript. C.D.B. designed experiments and wrote the manuscript. R.K.B., J.C.K., and K.P.L. conducted the experiments and assisted in data curation and analysis. M.I.D. assisted in the writing and editing of the manuscript. M.K.M. designed the bioinformatics pipeline and assisted in the analysis and writing of the manuscript

## DATA AVAILABILITY STATEMENT

All data in its raw and processed forms will be made immediately available upon request.

## SUPPLEMENTARY INFORMATION

### SUPPLEMENTAL METHODS

#### PF-CPP paradigm

On Day (**D**) 1, mice were assessed for initial preference for the PF-paired side (right side) over 30 min. On D2, D4, D9, D11, D16, and D18, mice were confined to the right side of the apparatus for 30 min and allowed access to PF. Food pellets were weighed both before and after each 30 min session. On D3, D5, D10, D12, D17, and D19, a clean, empty food bowl was provided in the corner on the left side and mice were confined there for 30 min. on D22, mice were provided with open access to both sides (clean, empty food bowls on both sides) and were re-assessed for time spent on the PF-paired side. PF-CPP was assessed by calculating the change in time (s) spent on the PF-paired side between D1 and D22 (D22-D1).

On D23, mice were tested in a light/dark conflict test for compulsive-like eating (**CLE**) (Kirkpatrick et al., 2017) in which PF was available in the light side of the box which is considered anxiety-provoking and aversive. PF Intake was calculated as % body weight consumed [g consumed / body weight (g) * 100]. All behaviors were recorded with infrared cameras (Swann Communications U.S.A. Inc., Santa Fe Springs, CA, USA) and video-tracked with ANY-maze video tracking software (Stoelting Co., Wood Dale, IL, USA).

#### High impact variants for GeneNetwork analyses

Ensembl-annotated “high impact variants” included the following classifications: “coding sequence variant,” “feature elongation,” “feature truncation,” “incomplete terminal codon variant,” “initiator codon variant,” “mature miRNA variant,” “missense variant,” “NMD transcript variant,” “regulatory region ablation,” “regulatory region amplification,” “regulatory region variant,” “splice acceptor variant,” “splice donor variant,” “splice region variant,” “stop gained,” “stop lost,” “TF binding site variant,” “TFBS ablation,” “TFBS amplification,” “transcript ablation,” or “transcript amplification.”

#### Genenetwork eQTL datasets from BXD RI strains

We used the following eQTL datasets for GeneNetwork (Mulligan et al., 2017) analyses: Amygdala mRNA [INIA Amygdala Cohort Affy MoGene 1.0 ST (Mar11); GN323], hypothalamus mRNA [INIA Hypothalamus Affy MoGene 1.0 ST (Nov10); GN281], Nucleus Accumbens mRNA [VCU BXD NAc Sal M430 2.0 (Oct07); GN156], striatum mRNA [HQF Striatum Affy Mouse Exon 1.0ST Gene Level (Dec09); GN399], Prefrontal Cortex mRNA [VCU BXD PFC Sal M430 2.0 (Dec06); GN135], and ventral tegmental area mRNA [VCU BXD VTA Sal M430 2.0 (Jun09); GN228].

### SUPPLEMENTARY RESULTS

In examining PF-CPP, we did not identify any genome-wide significant QTLs for the D22-D1 right side time measure (**Supplementary Figure 2**)., although we observed a significant female-selective QTL for initial preference for the PF-paired side (D1 right side time) that coincidentally, also happened to be located on chromosome 6 (but more distal to the BLE QTL) and drove a confounding, nonsignificant trending QTL at the same locus (**Supplementary Figure 2**). Thus, these results are not related to motivation for PF but rather, the confounding effect of the D1 QTL (initial preference, prior to PF exposure) on the D22-D1 subtraction measure.

### SUPPLEMENTARY TABLE LEGENDS

**Supplementary Table 1**. List of polymorphic genes, position (bp), variant name, allele, type, and consequence with Ensembl high-impact annotations within 1.5 LOD confidence interval for our male-specific chromosome 6 QTL (111.06 Mb – 124.76 Mb). Generated from Sanger Institute Mouse Genomes Project (https://www.sanger.ac.uk).

**Supplementary Table 2**. Phenotypes identified from GeneNetwork that have peak QTLs localized in the male-selective chromosome 6 QTL (111-125 Mb). Yellow-highlighted rows denote the phenotyped used in GeneNetwork analysis for *Adipor2, Plxnd1*, and *Rad18*.

### SUPPLEMENTARY FIGURE LEGENDS

**Supplementary Figure 1.**
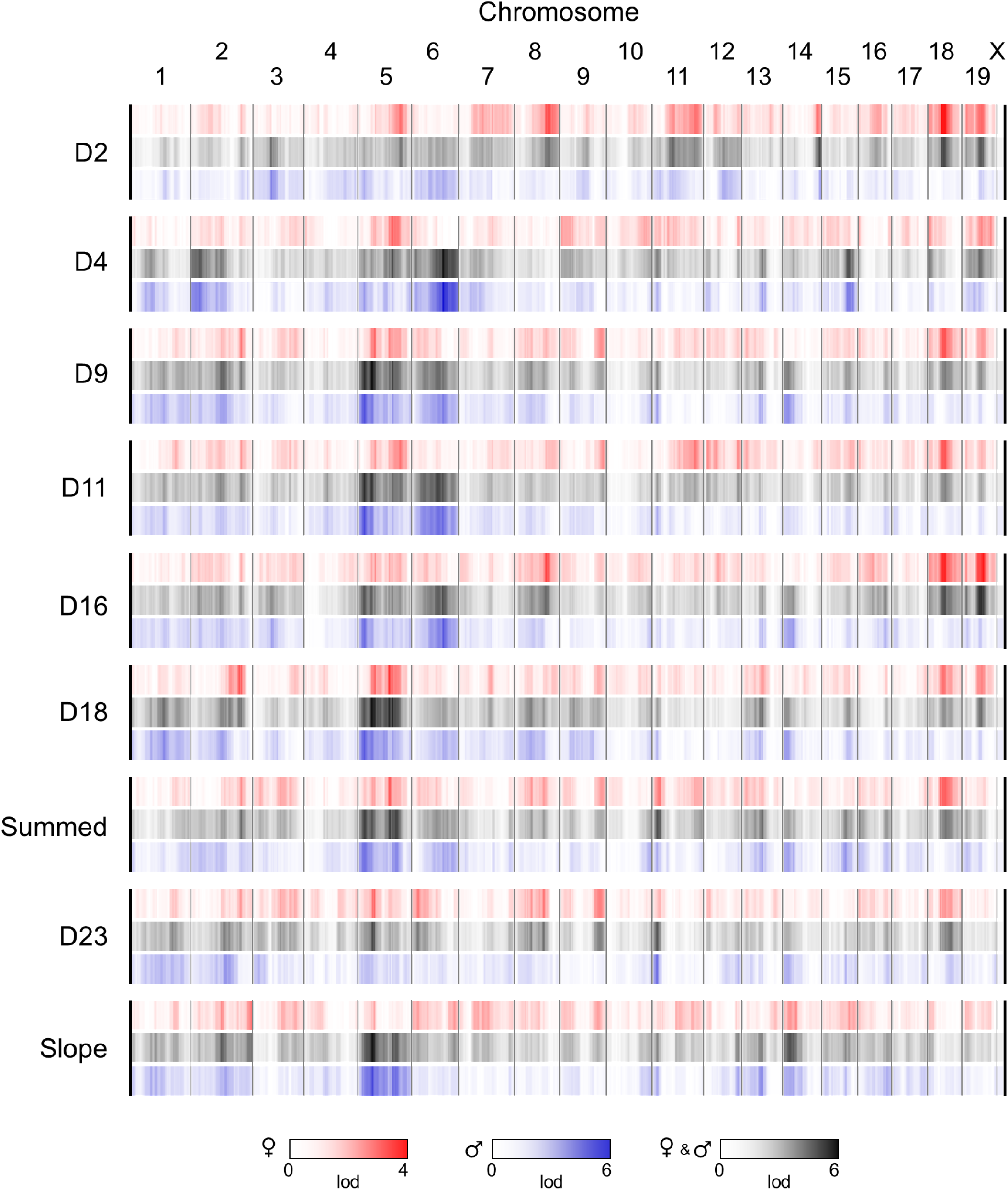
Linkage heat map of LOD scores at each marker for each day (D) of PF intake in B6J × D2J-F2 females (red), males (blue), and sex-combined (black) mice. The graph was generated using R/qtlcharts (Broman, 2015). QTLs for the sex-combined, females-only, and males-only analysis of PF intake across BLE training days and CLE test are presented, including the slope of escalation in PF intake across training days. In particular, note chromosomes 5 and 6 for the sex-combined and males-only analyses and chromosome 18 for the females-only analysis.

**Supplementary Figure 2.**
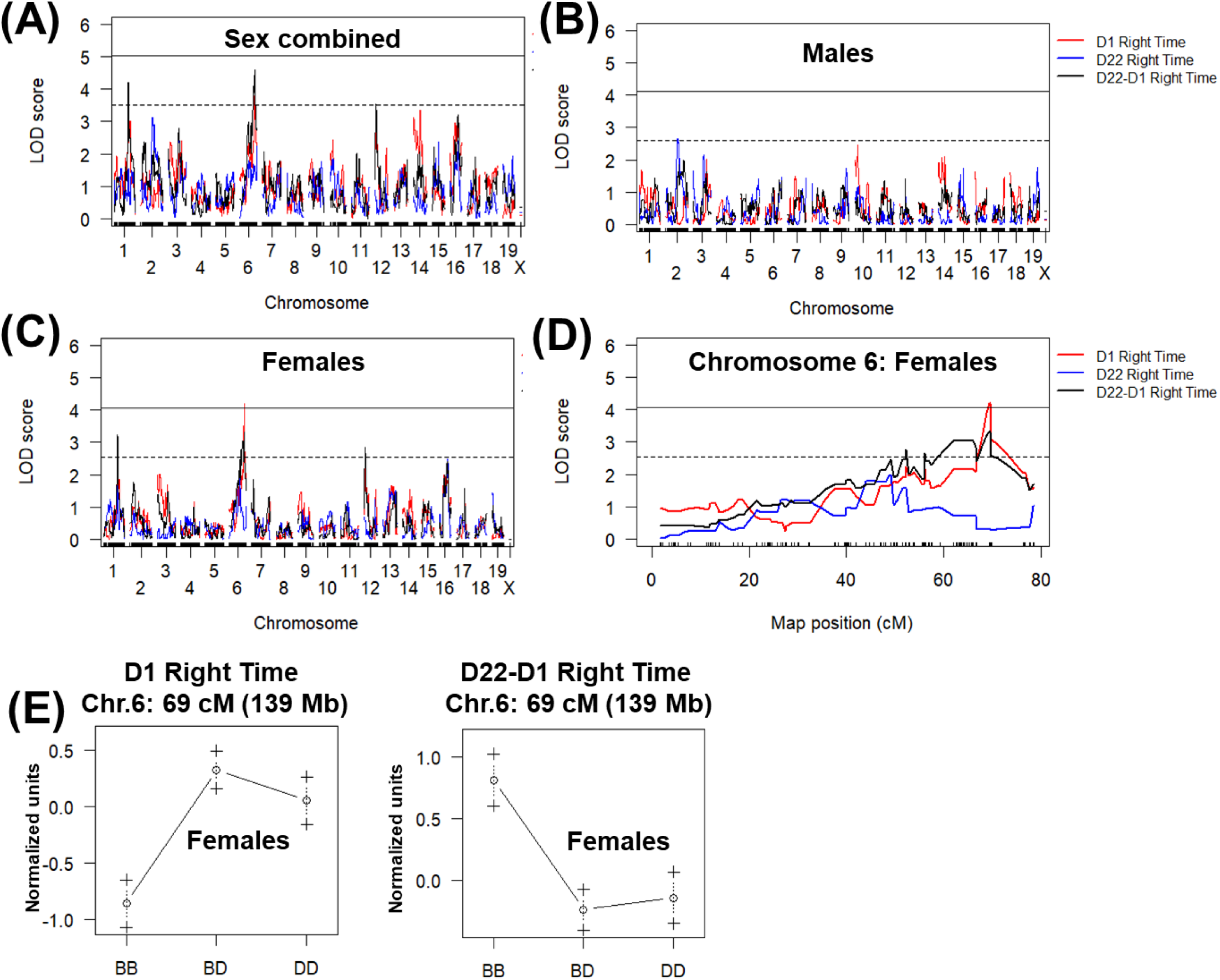
Genome-wide analysis PF-CPP. **(A-C):** Sex-combined, males-only, and females-only QTL analysis. A genome-wide significant QTL was identified in females (**C**) for initial preference for the PF-paired side (right side) on Day (D) 1. **(D):** Chromosome 6 QTL plot in females for D1 right side time and for D22-D1 right side time (Post-PF training). The D1 QTL interval peaked at 69 cM (139 Mb; LOD = 4.2; p < 0.05) and spanned 67-73 cM (137-142 Mb) or 52-77 cM (111-144 Mb; Bayes Credible interval). **(E):** Chromosome 6 effect plot of peak locus in females for the significant QTL (**left panel:** D1 time on PF-paired side; right side) and for the nonsignificant QTL (**right panel:** D22-D1 right side time).

**Supplementary Figure 3.**
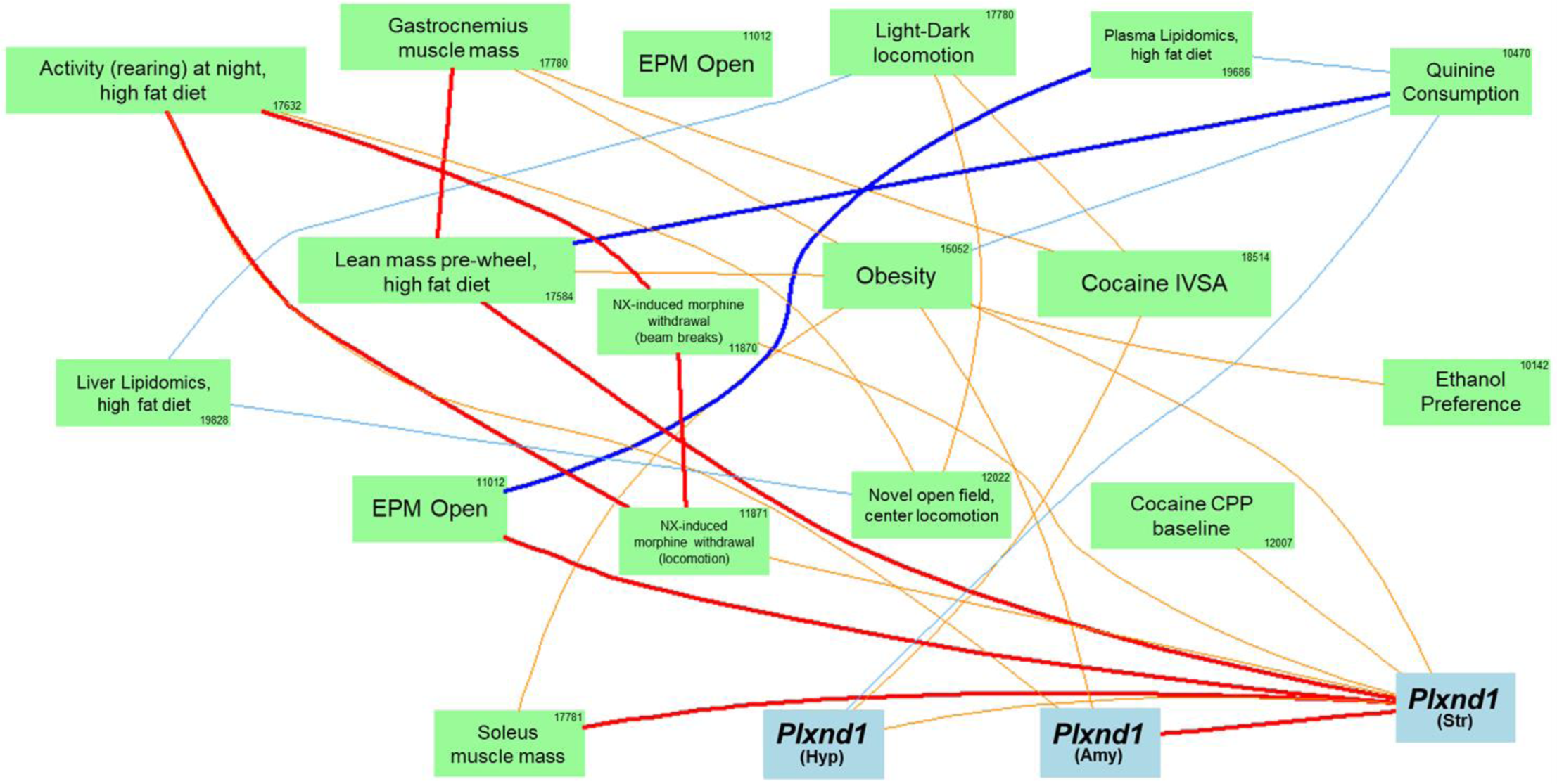
PheQTL-eQTL network graph for *Plxnd1* within the male-specific chromosome 6 QTL interval shows *cis-*eQTLs in three brain regions (hypothalamus, amygdala, striatum) with eight nodes. Correlated PheQTLs include traits related to body mass, drug addiction, obesity, and bitter taste. Bold blue lines indicate Pearson correlation coefficients of −1 to −0.7, normal light blue: −0.7 to −0.5, bold red: 0.7 to 1, normal orange: 0.5 to 0.7. The number inside each green box indicates the trait’s BXDPublish Record ID number. Only connected nodes are shown.

**Supplementary Figure 4.**
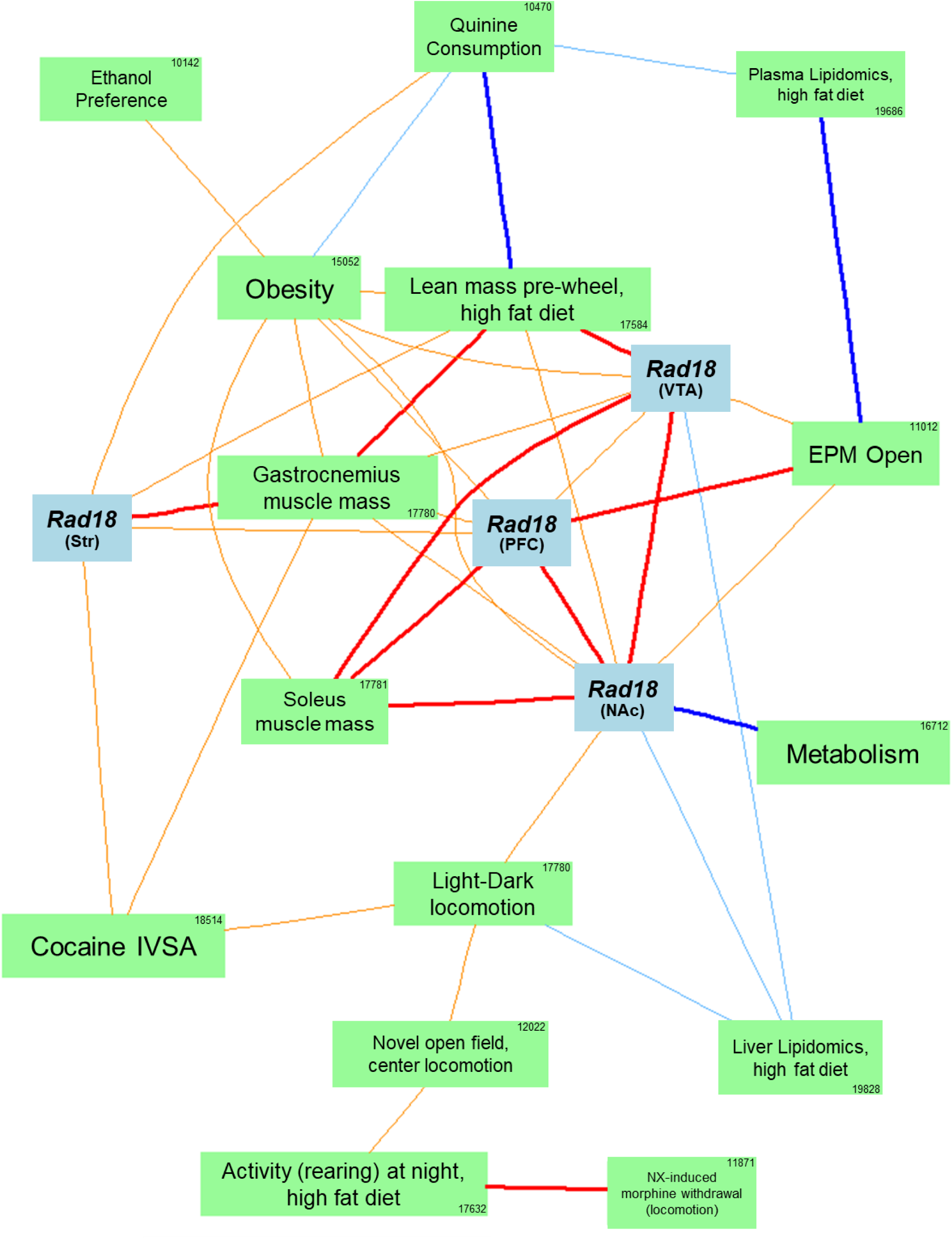
PheQTL-eQTL network graph for *Rad18* within the male-specific chromosome 6 QTL interval shows *cis-*eQTLs in three brain regions (hypothalamus, amygdala, striatum) with eight nodes. Correlated PheQTLs include traits involved in body mass, drug addiction, obesity, and bitter taste. Bold blue lines indicate Pearson correlation coefficients of −1 to −0.7, normal light blue: −0.7 to −0.5, bold red: 0.7 to 1, normal orange: 0.5 to 0.7. The number inside each green box indicates the trait’s BXDPublish Record ID number. Only connected nodes are shown.

